# Singe cell RNA sequencing data processing using cloud-based serverless computing

**DOI:** 10.1101/2025.04.26.650787

**Authors:** Ling-Hong Hung, Niharika Nasam, Chris Biju, Wes Lloyd, Ka Yee Yeung

## Abstract

Singe cell RNA sequencing (scRNA-seq) has become a routine method for measuring cell activities. Processing large scRNA-seq datasets requires high-performance computing resources. The emergence of cloud computing allows us to leverage its on-demand capabilities without major investment in infrastructure. Serverless computing provides cost efficiency by allowing users to pay only for actual resource usage, eliminating the necessity for pre-allocated server capacities. Additionally, there is no requirement to set up servers in advance. We present a novel and generalizable methodology using serverless cloud computing to accelerate computationally intensive workflows. We create an on-demand “supercomputer” using rapidly deployable cloud serverless functions as automatically provisioned computation units. We tested our methodology of optimizing a scRNA-seq workflow by leveraging serverless functions on the cloud using two publicly available peripheral blood mononuclear cell (PBMC) datasets. In addition, we demonstrate our approach using data generated by the NIH MorPhiC program, where we process a 450 GB human scRNA-seq dataset across 86 cell lines designed to study the temporal impact of perturbations on pancreatic differentiation. We compared the total execution time of the scRNA-seq serverless workflow with the traditional workflow without using serverless functions, and demonstrate major speedup for large scRNA-seq datasets.

## 1 Introduction

Singe cell RNA sequencing (scRNA-seq) measures the abundance of mRNA molecules at single cell resolution and has become a routine method in laboratories. The widespread adoption and technological advances of scRNA-seq have led to the development of many computational tools (21). An scRNA-seq data processing *workflow* typically consists of multiple steps, including data quality control, alignment of sequence fragments to a reference sequence, assignment of reads to genes and cells, cell barcode demultiplexing, UMI deduplication, and counting the number of unique RNA molecules. This procedure results in a gene-by-cell count matrix which is used as an estimate to quantify RNA molecules in each cell for each gene (27). Data normalization and subsequent downstream analysis are often performed using this gene-by-cell count matrix. Each step in an analytical workflow could have different requirements for computing resources. In particular, many methods and software tools have been developed to perform the computationally intensive alignment step, for example STARsolo (28), Cell Ranger (10), Piscem (19), and Alevin-fry(20).

Rapid scale-up of scRNA-seq technology (i.e. more cells and samples) coupled with the potential of combining with other data sources at single-cell resolution across modalities (e.g. spatial location) has established the crucial role of scalable computing resources. Cloud computing enables convenient, on-demand access to a shared pool of configurable computing resources that can be provisioned and released as needed. In particular, serverless computing adopts a simplified model to create scalable applications with reduced configuration and management overhead on the cloud (14). Instead of the user provisioning and managing virtual servers, the cloud provider automatically allocates machine resources as needed (13; 29). Most public cloud providers offer serverless computing capabilities. The pricing structure for serverless computing is based on actual application runtime, eliminating the cost of renting idle servers. However, serverless functions are designed for small microservices, with limited memory, disk space, and execution time. Here, we present a serverless pipeline for scRNA-seq, where the alignment step is executed through the invocation of multiple serverless functions on the cloud, and demonstrate reduction in execution time.

### 1.1 Related Work

Serverless computing is meant to let businesses and application developers focus on the program they need to run and not worry at all about the machine it is on or the resources it requires (29). In serverless computing, server infrastructure is created on demand and dedicated solely to the functions required to execute your code. Serverless computing offers cost savings through a pay-as-you-go model, eliminating the need for provisioning and maintaining dedicated servers. A primary advantage of serverless computing is its propensity for scalability and massive parallel computation. Serverless computing additionally benefits from leveraging flexible cloud services for data storage to provide on-demand access to applications and resources (15).

Grzesik et al. (17) reviewed serverless computing solutions in bioinformatics evaluating their usage in omics data analysis and integration. In their survey, they emphasize the utility of serverless computing for providing access to substantial computational resources while supporting the integration of diverse data sources within complex analysis pipelines. Their work highlights the acceleration potential provided by cloud computing, specifically through the serverless paradigm, which streamlines infrastructure management for developers. Their survey offers a critical review of serverless solutions in bioinformatics, while also describing their own application supporting multi-omics data analysis and integration. Notably, their evaluation extends to the effectiveness of these solutions within the context of the COVID-19 pandemic.

BioDepot Workflow Builder(Bwb) (24) is an open source cloud-enabled containerized platform that simplifies bioinformatics workflow creation through an interactive graphical interface, enabling biomedical researchers to design and customize complex analytical workflows. With drag-and-drop functionality and options to integrate with other open source software such as Jupyter, Fiji and QuPath (26; 30), Bwb streamlines the process of constructing and reproducibly executing complex bioinformatics workflows.

Hung et al. (25) introduced an innovative approach achieving a remarkable 1100-fold computational speedup in RNA sequencing analysis. This enabled biomedical scientists to dynamically fine-tune alignment parameters in real-time, facilitating iterative improvements in the final analytical results. They employed a publish and subscribe mechanism to invoke serverless instances to perform the compute intensive alignment step in which millions of short reads (sequences) are mapped to the reference sequence. Efficiency was achieved by splitting the input sequences into smaller pieces (or shards) and invoking more than 1700 serverless instances to perform alignment simultaneously on each data shard, thus resulting in reduced execution time in the alignment step in an RNA-seq workflow. In addition, Hung et al. implemented this serverless RNA-seq workflow using the Bwb platform to provide an interactive graphical interface for biomedical scientists (25).

Similar to the approach by Hung et al., Pietro et al .(15) leveraged serverless computing to support computation of genomic data, focusing on RNA-seq read-mapping to a reference genome, which is the most time-consuming task. Their experiments demonstrated a huge reduction in runtime when the read-mapping was performed on serverless instances, while demonstrating significant enhancements to scalability and parallelism possible with serverless computing. As another application of serverless techniques in bioinformatics, De Carvalho et al. proposed a framework to provision and execute protein sequence alignment workflows composed of multiple tasks (16). They compared the computation time of workflows executed using traditional virtual machines hosted on AWS EC2 instances with serverless computing using AWS Lambda functions. Their experiments confirmed that protein sequence alignments performed using Lambda functions executed in a shorter average time in relation to the alignment performed on EC2 instances.

While bulk RNA-seq provides estimates of the average expression level for each gene across a population of cells, scRNA-seq can estimate a distribution of expression levels for each gene across a population of cells (12). A typical scRNA-seq workflow contains the following steps: mapping the short sequences to a reference; assigning reads to genes and cells; and counting the number of unique RNA molecules. The output of these steps includes a gene/cell count matrix, and subsequent downstream analyses (such as visualization and clustering).

### 1.2 Our contributions

We present a novel and generalizable methodology using serverless cloud computing to accelerate computationally intensive workflows. Specifically, we present a serverless pipeline for scRNA-seq, where the alignment step is executed through the invocation of multiple serverless function instances. We optimized the Piscem-Alevin Fry pipeline (18) to surmount resource limitations of serverless functions by splitting large input sequence files into smaller chunks and using storage buckets to store input and output files. Each serverless function instance simultaneously maps reads from sequence input files to the reference. Our results demonstrate a major speedup for large scRNA-seq datasets due to the parallel alignment step. We have expanded the serverless approach previously employed by Hung et al. (25) in bulk RNA-seq to the realm of scRNA-seq.

## 2 Overview of Serverless scRNA-seq Workflow

Serverless functions are designed for small short-lived applications and accordingly, have resource limitations. As an example, Amazon Web Services (AWS) Lambda functions are limited to 15 minutes maximum runtime, and up to 10 GB of memory (6). In particular, we developed an optimized cloud-based serverless scRNA-seq workflow by leveraging the Piscem-Alevin Fry pipeline (18) that is accurate, scalable, fast, and memory-efficient. Piscem’s process of mapping reads to the reference is considerably faster than conventional aligners, while Alevin-Fry (20) is designed for rapid processing of barcodes and UMIs with minimal memory usage. These features make the pipeline ideal for large-scale scRNA-seq studies, particularly in cloud-based serverless environments where computational and memory efficiency are critical. See the Methods section for a detailed summary of the Piscem-Alevin Fry pipeline (18).

Our serverless scRNA-seq workflow is implemented using the serverless AWS Lambda function, and the overall architecture is illustrated in Figure 1. The pipeline starts with an initialization step (steps 1, 2, 3 in Figure 1) in which AWS resources are created and input FASTQ files are pre-processed. Due to the small and transient nature of the disk space allotted to a function, a storage bucket (AWS Simple Storage Service S3) is used to store input files that need to be transferred to a function, and output files that are generated by the function that need to be returned to the client. The bucket provides an intermediary for data transfer between the client and the serverless function instances, assuming the role of a distributed file system for a traditional “supercomputer”.

**Figure 1:**
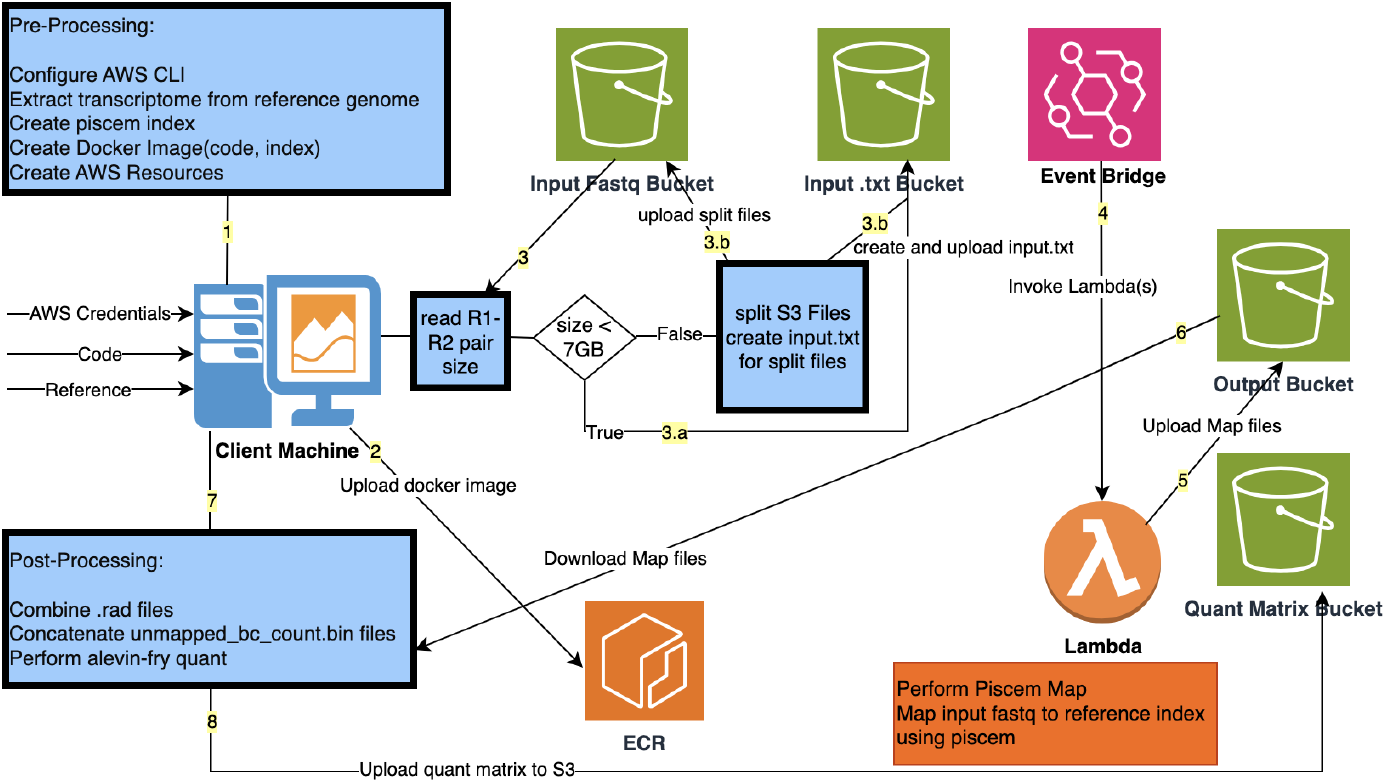
Overview of our optimized Piscem Alevin-Fry serverless architecture. Steps 1, 2 and 3 represent the pre-processing stage where AWS resources are created and input sequence files are split if the sequence file size exceeds 7 GB. Steps 4 and 5 represent serverless execution. Steps 6 and 7 represent the post-processing phase in which output files are downloaded from S3 and subsequently merged before the Alevin-Fry step is run. In step 8, output files from Alevin quant are uploaded to S3.

One of our design challenges is to modify the alignment process so it can execute within the limitations of serverless function instances while simultaneously reducing and mitigating the effect of the data transfers between the client, bucket, and serverless layers. This was accomplished by reducing the size of input FASTQ files if necessary. After the reference and executables are downloaded to the serverless instance, there is approximately 7 GB of space remaining for the input FASTQ files. If the file size exceeds 7GB, the input files are split into smaller chunks and uploaded to S3. A counter is maintained to track the number of uploaded input files. See steps 2 and 3 in Figure 1.

The alignment step in which reads are mapped to the reference is performed using multiple instances of an ‘alignment’ serverless Lambda function. For large input FASTQ files (i.e. greater than 7 GB), multiple Lambda functions are executed simultaneously. Each Lambda function downloads the input (potentially split) files from the corresponding S3 bucket. Once downloaded, `piscem map` (19) is executed, aligning the reads to the reference genome. The resulting output files are then uploaded to the designated output S3 bucket. See steps 4 and 5 in Figure 1.

After all output files from the alignment step have been generated, the processed output files are downloaded to the client machine (step 6 in Figure 1). These output files are subsequently merged and then processed using Alevin-Fry which identifies valid cell barcodes, performs barcode correction, and barcode demultiplexing. Finally, `alevin-fry quant` (9) is executed to perform UMI deduplication and generate the gene expression count matrix (step 7 in Figure 1).

The final quantification matrix is uploaded to an S3 bucket. All AWS resources created during the pipeline setup phase are deleted. Additionally, any split files uploaded to the input FASTQ bucket during processing are removed. See step 8 in Figure 1.

## 3 Data Description

We tested our serverless scRNA-seq workflow using three publicly available datasets. Our initial test cases for pipeline validation include two peripheral blood mononuclear cells (PBMCs) datasets provided by 10x Genomics. The first dataset (denoted as **PBMC 1K**) consists of 1,222 PBMCs from a healthy donor, with an approximate size of 5 GB. It is publicly available (2). The second dataset (denoted as **PBMC 10K**) contains 11,769 PBMCs from a healthy donor, with a larger size of approximately 45 GB. It is publicly available for download (1).

Following the validation phase, a comprehensive analysis is conducted using a dataset generated by the Memorial Sloan Kettering Cancer Center (MSKCC) as part of the NIH MorPhiC (Molecular Phenotypes of Null Alleles in Cells) program (11). This scRNA-seq dataset (denoted as **KO dataset**) is designed to study the temporal impact of perturbations on pancreatic differentiation. Specifically, it includes knockout (KO) data spanning 86 cell lines across five distinct time points. The total dataset size is approximately 450 GB. This dataset is publicly available from the NCBI Gene Expression Omnibus (GEO) with accession number GSE288587 (22) and also at the European Nucleotide Archive (ENA) with accession number PRJEB85005 (23).

The PBMC datasets was used to validate the accuracy of the serverless pipeline. Benchmarking and performance evaluation of the pipeline are performed using the MorPhiC knockout dataset.

## 4 Experimental Results

This section presents the results of the serverless execution times across different datasets and compares the performance against an on-server execution approach. The datasets used include the **KO dataset, PBMC 1K**, and **PBMC 10K**. The execution time for each processing step is recorded and analyzed.

### 4.1 Experimental Setup

The serverless pipeline is invoked from an AWS m6id.16xlarge EC2 instance, which has 64 vCPUs, 256 GiB RAM, up to 50 Gbps networking, and 3.8 TB NVMe SSD storage, optimized for compute-intensive workloads.

For the traditional (non-serverless) pipeline, it is run on a dedicated server equipped with a 13th Gen Intel Core i9-13900KF processor, 125 GB RAM, 16.4 TB HDD (RAID1), and 1.8 TB NVMe SSD (RAID1), running Ubuntu 22.04.2 LTS.

#### Account-dependent AWS limits

Reproducing the serverless benchmarks may require AWS quota adjustments that vary by account and region. In some AWS accounts, the maximum configurable AWS Lambda memory (and thus the available vCPU allocation) can be lower than the configuration used in this study (e.g., capped at 3008 MB, corresponding to fewer vCPUs). Under such limits, the `piscem map` thread count must be reduced (e.g., `-t 2` instead of `-t 6`), and the mapping step can be approximately three times slower, while producing the same output files. If higher Lambda memory is not available, one mitigation is to increase parallelism by splitting large inputs into more shards (i.e., more parts), invoking more concurrent alignment functions, and then merging the resulting RAD outputs on the client. Because AWS Lambda billing scales with memory × runtime, this approach typically yields similar total cost while improving wall-clock throughput. Users may also need to request an EC2 on-demand vCPU quota increase (or choose an equivalent instance type) to launch the orchestration instance class used in this study.

### 4.2 Serverless Execution Time

Table 1 shows the execution times for different tasks in the serverless pipeline. The overall processing time in the bottom row includes pre-processing, execution, and post-processing steps. Note that serverless Lambda function instances are being triggered with execution initiated during the “split and upload” step, while the “piscem map” step represents the additional time required to complete the alignment step. The parallel execution of multiple Lambda function instances results in the reduction of the execution step in the “piscem map” step for the large KO dataset (0.52 minutes). For the PBMC 1K dataset, the input sequence files are less than 7 GB, so there was no splitting of input files, and the “piscem map” step is performed using a single Lambda function call. On the other hand, the input sequence files are more than 7 GB for the PBMC 10K dataset, so the input files were split and multiple Lambda function instances were triggered and executed in parallel. This is the reason why the execution for the “piscem map” step for the PBMC 10K dataset is smaller than that of the PBMC 1K dataset (1.51 minutes for PBMC 10K vs. 3.51 minutes for PBMC 1K)

**Table 1:** Execution time (in minutes) applying our serverless scRNA-seq pipeline to the knockout (KO), PBMC 1k and PBMC 10k datasets. The corresponding step number in Figure 1 is shown in column 2. Note that these execution times do not include the amount of time fetching the raw sequence data before the “split and upload” step.

| Task | Step # | KO | PBMC 1K | PBMC 10K |
| --- | --- | --- | --- | --- |
| Files Size |  | 453.43 GB | 4.69GB | 44.05GB |
| Split and Upload<br>[on-server] | 3 | 14.15 | 0.11 | 8.84 |
| Piscem Map<br>[serverless] | 4 | 0.52 | 3.51 | 1.51 |
| Download Files<br>[on-server] | 6 | 1.81 | 0.06 | 0.30 |
| Total Processing |  | 17.49 | 4.68 | 11.65 |
| Combined .rad<br>[on-server] | 7 | 0.82 | 0.00 | 0.10 |
| Concatenate un-<br>mapped_bc_count.bin<br>[on-server] | 7 | 0.01 | 0.00 | 0.00 |
| Alevin [on-server] | 7 | 11.58 | 0.10 | 0.88 |
| Upload Output<br>Files [on-server] | 8 | 0.56 | 0.01 | 0.05 |
| **Overall Process** |  | **30.46** | **4.80** | **12.68** |

### 4.3 Serverless vs. On-Server Comparison

Table 2 compares the execution time between the serverless pipeline and traditional on-server processing of the KO, PBMC 1K, and PBMC 10K scRNA-seq datasets. The “serverless execution” time represents the “overall processing” time from Table 1. “Speedup” is calculated by the ratio of the on-server execution time to the serverless execution time. Note that the serverless scRNA-seq pipeline achieved a speed up of 1.80x and 6.45x for the PBMC 10K and KO datasets respectively. In contrast, the serverless execution time more than doubled that of the on-server processing of the smaller PBMC 1K dataset. The serverless workflow is highly efficient for larger datasets due to the parallel execution of multiple serverless function instances over the split input files. On the other hand, there is minimal or no performance gain for small datasets due to time spent on creating cloud resources and infrastructure overhead.

**Table 2:** Execution time (in minutes) comparing our serverless versus on-Server processing of the knockout (KO), PBMC 1k and PBMC 10k datasets.

| Method | KO | PBMC 1K | PBMC 10K |
| --- | --- | --- | --- |
| Files Size | 453.43 GB | 4.69 GB | 44.05 GB |
| Serverless Execution | 30.46 | 4.80 | 12.86 |
| On-Server (SSD) Execution | 196.38 | 2.26 | 23.21 |
| **Speedup** | **6.45x** | **0.47x** | **1.80x** |

## 5 Discussion

We have demonstrated how serverless cloud instances can be harnessed to provide on-demand and scalable “supercomputing” without the need for specialized hardware or a large cluster of server nodes. Our experimental results show that our serverless approach significantly reduces execution time for larger datasets (e.g., KO dataset). The parallel processing capabilities of AWS Lambda allow for efficient scaling. For smaller datasets such as PBMC 1K, serverless execution introduces an overhead, making it less efficient compared to an on-server SSD execution. Our serverless approach achieves up to 6.44x speedup for the large KO dataset. However, for small datasets, the additional overhead of initializing serverless infrastructure resulted in negative performance improvement. These findings suggest that serverless workflows are particularly useful for processing large-scale genomic data, but may not be suitable for small datasets where traditional computing methods offer better performance.

## 6 Methods

### 6.1 Steps Involved in Single-Cell RNA Sequencing

Several computational steps are generally involved in processing raw scRNA-seq data into a gene-by-cell count matrix. In the *transcriptome annotation step*, the reference is annotated using a gene annotation file, which provides the locations of genes and transcripts. This step is crucial for accurately mapping sequencing reads to known genes. The *transcriptome indexing* step facilitates rapid and efficient read alignment by creating an index for the annotated transcriptome. Indexing allows alignment tools to quickly search and match sequencing reads to reference sequences. In the *read alignment step*, sequenced reads are aligned to the indexed reference transcriptome. This step assigns each read to its most likely transcript, enabling gene-level quantification. Next, barcodes are filtered based on the cumulative frequency distribution of reads associated with each barcode. This *permit list generation* step ensures that only valid barcodes representing real cells are retained. In the *barcode correction* step, sequencing errors in cell barcodes are corrected by matching them to the closest accepted barcodes. This improves data accuracy by reducing barcode mismatches. Next, reads are grouped according to their corrected cell barcodes, assigning each read to its respective single cell. This *barcode demultiplexing* step enables downstream cell-specific analysis. Unique Molecular Identifiers (UMIs) are used to remove duplicate reads that share the same barcode, UMI, and gene combination. This *UMI deduplication* step prevents amplification bias from affecting gene quantification. Finally, a gene expression count matrix is generated, where rows correspond to genes, columns correspond to cells, and values represent the number of times a gene is detected in a given cell. This matrix serves as the foundation for downstream analyses such as clustering and differential expression analysis.

### 6.2 Piscem-Alevin Fry Pipeline

The Piscem-Alevin Fry pipeline (18) is a highly efficient and scalable solution for processing scRNA-seq data. Alevin-fry is not only faster and more memory-efficient than other accurate quantification methods, but it also improves memory scalability and reduces false-positive expression issues found in other lightweight tools (20). Piscem is a lightweight mapping tool optimized for speed and memory efficiency, making it well suited for cloud-based serverless execution. Alevin-Fry (20), a companion tool, is designed to efficiently handle barcode correction, UMI deduplication, and count matrix generation, further enhancing the overall performance of the workflow (18).

The pipeline takes as input raw sequencing data in the form of FASTQ files, along with a reference genome file. The workflow begins with transcriptome annotation, where genes are mapped onto the reference genome to establish a structured representation of gene locations. Next, a transcript-to-gene mapping file is generated, providing the necessary reference for downstream analysis.

The first computational step in the pipeline involves indexing the transcriptome using the command `piscem build`. This indexing step is crucial for accelerating the alignment process in subsequent steps. Once the index is prepared, raw sequencing reads are mapped to the reference transcriptome using `piscem map`, a highly optimized mapping algorithm that efficiently aligns reads while maintaining accuracy.

Following the mapping step, Alevin-Fry is used to process and refine the data. The first step, executed via `alevin-fry generate-permit-list` (8), involves generating a permit list, which identifies valid cell barcodes from the sequencing data. This is followed by the `alevin-fry collate` (7) step, where barcode correction is performed to adjust for sequencing errors, and barcode demultiplexing is applied to assign reads to individual cells.

The final step in the pipeline is gene quantification. The `alevin-fry quant` (9) command performs UMI deduplication, ensuring that duplicate reads originating from PCR amplification are removed. This step also generates the final count matrix, where each row represents a gene, each column represents a single cell, and each value corresponds to the number of times a gene was detected in that specific cell.

### 6.3 Implementation: Serverless scRNA-seq Workflow on the Cloud

The pipeline begins with the input parameters, which include Amazon Web Services (AWS) credentials, code, and a reference genome file. The input FASTQ files are stored in an S3 (5) bucket, labeled as **Input FASTQ Bucket** in Figure 1. The first step in the workflow involves configuring the AWS Command Line Interface (CLI) on the client machine. Once configured, the transcriptome is annotated from the reference genome, followed by the generation of a transcript-to-gene mapping file. The annotated transcriptome is then indexed using `piscem index`. The credentials file generated after AWS CLI configuration, along with the transcriptome index, is bundled with the pipeline’s code, and a Docker image is created.

#### 6.3.1 Creation of Cloud Resources

The next step involves creating the necessary AWS resources. An Amazon Elastic Container Registry (ECR) (3) repository is created to store the Docker image. The image is built locally and then pushed to the ECR repository. This is labeled as step 1 in Figure 1. A Lambda (6) execution role is defined with the required policies to grant access to S3 resources. Subsequently, a Lambda function is created and configured to execute containerized workloads. Several S3 buckets are also set up: the **output bucket** to store Lambda-generated outputs, the **input** .**txt files bucket** to hold generated `input.txt` files, and the **final output bucket** to store the final Alevin-Fry quantification results. Additionally, an EventBridge rule is created to trigger the Lambda function upon new file uploads to the input `.txt` bucket, and appropriate permissions are granted.

Once AWS resources are configured, the client machine reads each R1-R2 file pair size from S3. If the file pair size is less than 7GB, an `input.txt` file is generated, containing the S3 paths of the corresponding R1-R2 file pairs, and is uploaded to the input `.txt` files bucket. If the file size exceeds 7GB, the file pairs are split into smaller chunks and uploaded to S3. For each split file, a corresponding `input.txt` file is created, containing the paths of the smaller file pairs, and uploaded to S3. To maintain uniqueness, each `input.txt` filename includes extracted lane and part information. A counter is maintained to track the number of uploaded input files. These are labeled as steps 2 and 3 in Figure 1.

#### 6.3.2 Lambda Configuration and Execution

The Lambda function in this pipeline is container-based and is configured to pull the image stored in the ECR repository. The function is allocated 10GB of memory and 10GB of internal storage. An EventBridge(4) rule is defined to trigger Lambda execution upon the upload of an `input.txt` file to the designated S3 bucket.

Upon execution, the Lambda function reads the `input.txt` file and downloads the R1 and R2 files from the S3 paths specified in the file. These files are provided as input parameters during the EventBridge-triggered Lambda execution. Once downloaded, `piscem map` (19) is executed, aligning the reads to the reference genome. The resulting output files are then uploaded to the designated output S3 bucket. To ensure unique output directory names, each output folder is named using the extracted lane and part of the information from the input file pairs. These are labeled as steps 4 and 5 in Figure 1.

#### 6.3.3 Post-Processing and Quantification

After all input files have been processed and uploaded, the client machine continuously polls the output S3 bucket to monitor the number of output folders. Once the count of input folders matches the count of output folders, the processed output files are downloaded to the client machine. This is labeled as step 6 in Figure 1. Each output folder contains `map.rad` and `unmapped_bc_count.bin` files. These files are merged using the `radtk` utility, resulting in a single concatenated `.rad` file and a single `.bin` file.

The merged files are then processed using Alevin-Fry. The first step involves running `alevin-fry generate-permit-list` (8), which identifies valid cell barcodes based on the cumulative frequency distribution of reads associated with each barcode. Next, `alevin-fry collate` (7) is executed to perform barcode correction and barcode demultiplexing. Finally, `alevin-fry quant` (9) is executed to perform UMI deduplication and generate the gene expression count matrix. This is labeled as step 7 in Figure 1.

The final quantification matrix is uploaded to the Quant Matrix S3 bucket to store final results. After this step, all AWS resources created during the pipeline setup phase are deleted. Additionally, any split files uploaded to the input FASTQ bucket during processing are removed, restoring the input bucket to its original state before pipeline execution. This is labeled as step 8 in Figure 1.

## 7 Data availability

The knockout (KO) datasets supporting the results of this article are available in the NCBI Gene Expression Omnibus (GEO) repository with accession number GSE288587 (22) and also at the European Nucleotide Archive (ENA) with accession number PRJEB85005 (23). The **PBMC 1K** data are publicly available at https://www.10xgenomics.com/datasets/1-k-pbm-cs-from-a-healthy-donor-v-3-chemistry-3-standard-3-0-0 The **PBMC 10K** data are publicly available at https://www.10xgenomics.com/datasets/10-k-pbm-cs-from-a-healthy-donor-v-3-chemistry-3-standard-3-0-0.

## 8 Competing Interests

L.H.H. and K.Y.Y. have equity interest in Biodepot LLC. The terms of this arrangement have been reviewed and approved by the University of Washington in accordance with its policies governing outside work and financial conflicts of interest in research.

## 9 Funding

L.H.H., N.N. and K.Y.Y. are supported by the National Institutes of Health (NIH) grant U24HG012674. N.N., C.B. and K.Y.Y. are also supported by the Virginia and Prentice Bloedel Endowment at the University of Washington.

## 10 Author’s Contributions

L.H.H. and K.Y.Y. conceived and supervised the study. N.N. developed and tested the tool and performed the analysis. L.H.H. designed the serverless scRNA-seq pipeline and empirical studies. N.N. and K.Y.Y. wrote the draft manuscript. C.B. assisted in testing and documentation. L.H.H., W.L. and K.Y.Y. were involved in discussions and contributed to the manuscript. All authors read and approved the final manuscript.

## 11 Acknowledgements

We would like to thank the Danwei Huangfu Lab at the Memorial Sloan Kettering Cancer Center and NIH grant UM1HG012654 used for generating the MorPhiC KO dataset. We would like to thank the data ingestion team at the European Bioinformatics Institute (Anu Shivalikanjli, Galabina Yordanova, Alexandros Orges Koci, Sandeep Selvakumar) for data collection, metadata annotation and brokering of the MorPhiC KO dataset.

## Notes

### Summary of Updates

Added Chris Biju as a co-author, who assisted in testing and checking correctness of results. He also updated software and corresponding documentation. Added a paragraph called "Account dependent AWS Limits" under Section 4.1 "Experimental Setup" to document account setup required to run our scripts.

## References

[1] 10k pbmcs from a healthy donor. https://www.10xgenomics.com/datasets/10-k-pbm-cs-from-a-healthy-donor-v-3-chemistry-3-standard-3-0-0.

[2] 1k pbmcs from a healthy donor. https://www.10xgenomics.com/datasets/1-k-pbm-cs-from-a-healthy-donor-v-3-chemistry-3-standard-3-0-0.

[3] Amazon elastic container registry (ecr). https://aws.amazon.com/ecr/.

[4] Amazon eventbridge. https://aws.amazon.com/eventbridge/.

[5] Amazon simple storage service (s3). https://aws.amazon.com/s3/.

[6] Aws lambda. https://aws.amazon.com/lambda/.

[7] Collate. https://alevin-fry.readthedocs.io/en/latest/collate.html.

[8] Generating permit list. https://alevin-fry.readthedocs.io/en/latest/generate_permit_list.html.

[9] Quant. https://alevin-fry.readthedocs.io/en/latest/quant.html.

[10] What is cell ranger? https://support.10xgenomics.com/single-cell-gene-expression/software/pipelines/latest/what-is-cell-ranger.

[11] Mazhar Adli, Laralynne Przybyla, Tony Burdett, Paul W Burridge, Pilar Cacheiro, Howard Y Chang, Jesse M Engreitz, Luke A Gilbert, William J Greenleaf, Li Hsu, et al. MorPhiC Consortium: towards functional characterization of all human genes. Nature, 638(8050):351–359, 2025.

[12] Tallulah S Andrews, Vladimir Yu Kiselev, Davis McCarthy, and Martin Hemberg. Tutorial: guidelines for the computational analysis of single-cell rna sequencing data. Nature protocols, 16(1):1–9, 2021.

[13] Ioana Baldini, Paul Castro, Kerry Chang, Perry Cheng, Stephen Fink, Vatche Ishakian, Nick Mitchell, Vinod Muthusamy, Rodric Rabbah, Aleksander Slominski, et al. Serverless computing: Current trends and open problems. Research advances in cloud computing, pp. 1–20, 2017.

[14] Paul Castro, Vatche Ishakian, Vinod Muthusamy, and Aleksander Slominski. The server is dead, long live the server: Rise of serverless computing, overview of current state and future trends in research and industry. arXiv preprint arXiv:1906.02888, 2019.

[15] Pietro Cinaglia, Jose Luis Vazquez-Poletti, and Mario Cannataro. Massive parallel alignment of rna-seq reads in serverless computing. Big Data and Cognitive Computing, 7(2):98, 2023.

[16] Leonardo Reboucas de Carvalho, Alba Cristina Alves Melo, and Aleteia Araujo. A framework for executing protein sequence alignment in cloud computing services. In Anais do XXII Simposio em Sistemas Computacionais de Alto Desempenho, pp. 48–59. SBC, 2021.

[17] Piotr Grzesik, Dariusz R Augustyn, L ukasz Wycislik, and Dariusz Mrozek. Serverless computing in omics data analysis and integration. Briefings in bioinformatics, 23(1):bbab349, 2022.

[18] Dongze He and Rob Patro. simpleaf: a simple, flexible, and scalable framework for single-cell data processing using alevin-fry. Bioinformatics, 39(10):btad614, 2023.

[19] Dongze He, Charlotte Soneson, and Rob Patro. Understanding and evaluating ambiguity in single-cell and single-nucleus rna-sequencing. bioRxiv 2023.01.04.522742, 2023.

[20] Dongze He, Mohsen Zakeri, Hirak Sarkar, Charlotte Soneson, Avi Srivastava, and Rob Patro. Alevin-fry unlocks rapid, accurate and memory-frugal quantification of single-cell rna-seq data. Nature Methods, 19(3):316–322, 2022.

[21] Lukas Heumos, Anna C Schaar, Christopher Lance, Anastasia Litinetskaya, Felix Drost, Luke Zappia, Malte D Lucken, Daniel C Strobl, Juan Henao, Fabiola Curion, et al. Best practices for single-cell analysis across modalities. Nature Reviews Genetics, 24(8):550–572, 2023.

[22] Danwei Huangfu, Dazhi Liu, Dapeng Yang, and Renhe Luo. Mapping the temporal impact of perturbations on lineage plasticity and gene regulation. https://www.ncbi.nlm.nih.gov/geo/query/acc.cgi?acc=GSE288587.

[23] Danwei Huangfu, Dazhi Liu, Dapeng Yang, and Renhe Luo. Morphic msk pooled scrna seq 202401. https://www.ebi.ac.uk/ena/browser/view/PRJEB85005.

[24] Ling-Hong Hung, Jiaming Hu, Trevor Meiss, Alyssa Ingersoll, Wes Lloyd, Daniel Kristiyanto, Yuguang Xiong, Eric Sobie, and Ka Yee Yeung. Building containerized workflows using the biodepot-workflow-builder. Cell systems, 9(5):508–514, 2019.

[25] Ling-Hong Hung, Xingzhi Niu, Wes Lloyd, and Ka Yee Yeung. Accessible and interactive rna sequencing analysis using serverless computing. bioRxiv 576199, 2020.

[26] Ling-Hong Hung, Evan Straw, Shishir Reddy, Robert Schmitz, Zachary Colburn, and Ka Yee Yeung. Cloud-enabled biodepot workflow builder integrates image processing using fiji with reproducible data analysis using jupyter notebooks. Scientific Reports, 12(1):14920, 2022.

[27] Byungjin Hwang, Ji Hyun Lee, and Duhee Bang. Single-cell rna sequencing technologies and bioinformatics pipelines. Experimental & molecular medicine, 50(8):1–14, 2018.

[28] Benjamin Kaminow, Dinar Yunusov, and Alexander Dobin. Starsolo: accurate, fast and versatile mapping/quantification of single-cell and singlenucleus rna-seq data. bioRxiv 2021.05.05.442755, 2021.

[29] Neil Savage. Going serverless. Communications of the ACM, 61(2):15–16, 2018.

[30] Pritpal Singh, Jocelyn H. Wright, Kimberly S. Smythe, Bryce Fukuda, Ling-Hong Hung, Cecilia CS Yeung, and Ka Yee Yeung. Graphical and interactive spatial proteomics image analysis workflow. bioRxiv, 2025.

